# Cloacal virome of an ancient host lineage – the tuatara (*Sphenodon punctatus*) – reveals abundant and diverse diet-related viruses

**DOI:** 10.1101/2022.06.14.496210

**Authors:** Stephanie J. Waller, Sarah Lamar, Benjamin J. Perry, Rebecca M. Grimwood, Edward C. Holmes, Jemma L. Geoghegan

## Abstract

Tuatara (*Sphenodon punctatus*) are one of the most phylogenetically isolated species and provide a unique host system to study virus evolution. While the tuatara genome, sequenced in 2020, revealed many endogenous viral elements, we know little of the exogenous viruses that infect tuatara. We performed a metatranscriptomics study of tuatara cloaca samples from a wild population on Takapourewa (Stephens Island), Aotearoa New Zealand. From these data we identified 49 potentially novel viral species that spanned 20 RNA viral families and/or orders, the vast majority (48) of which were likely dietary related. Notably, using a protein structure homology search, we identified a highly divergent novel virus within the *Picornaviridae* which may directly infect tuatara. Additionally, two endogenous tuatara adintoviruses were characterised that exhibited long-term viral-host co-divergence. Overall, our results indicate that the tuatara cloacal virome is highly diverse likely due a large number of dietary related viruses.

## 2. Introduction

Tuatara (*Sphenodon punctatus*) are endemic to Aotearoa New Zealand and are the sole surviving member of the once highly diverse and widely distributed reptilian order Rhynchocephalia (Sphenodontida) (1). Rhynchocephalians diverged from the Squamata order (lizards, snakes and amphisbaenians) more than 250 million years ago (2) and are an important evolutionary link to the now extinct stem reptiles from which modern vertebrates evolved (1,3). The tuatara genome was sequenced in 2020 (3), within which more than 450 endogenous tuatara retroviruses have been identified (3–6). However, little is known about the exogenous viruses that infect these hosts, although such information provides an opportunity to better understand how vertebrate viruses have evolved across the reptilian order.

Since tuatara have evolved in isolation for millions of years they provide a unique window on virus evolution. Zealandia, the continental crust from which New Zealand was formed, split from Gondwana around c. 84 million years ago (7). Following this event, the country’s flora and fauna evolved in near isolation from the rest of the world resulting in high levels of endemism (8). Fossil records indicate that tuatara once lived throughout mainland New Zealand (1). Following the arrival of the first humans ~800 years ago, tuatara populations rapidly declined due to habitat loss, the introduction of mammalian predators (1), and rising global temperatures that have affected sex ratios (9). Tuatara can now be found naturally on approximately 32 off-shore islands (10). While the predator-free island of Takapourewa (Stephens Island) supports the largest naturally occurring population of tuatara, estimated to be between 30,000-50,000 (1), many of the other offshore island populations consist of an estimated 10-100 tuatara which are extremely vulnerable to threats such as disease (10). Indeed, tuatara have been categorised as an ‘At Risk, Relict’ species that have experienced a rapid decline in numbers over the past several centuries and occupy less than 10% of their former habitat range (11).

Although viral infection poses an obvious threat to the health and survival of tuatara, our current knowledge of the tuatara virome is extremely limited (3–6). Revealing the viruses that infect tuatara could assist in future conservation efforts to protect these *taonga* (treasured) species. Here, we uncovered the faecal virome of the tuatara. In addition, we aimed to assess whether the health of the tuatara, measured by body condition index (BCI), a mass to length-based ratio, influenced viral diversity, richness and abundance (12).

## 3. Materials and Methods

### 3.1 Animal Ethics Approval

This project was approved by the Victoria University of Wellington Animal Ethics Committee (#27041) and the Department of Conservation (Wildlife Authority Act Number 50568 - FAU).

### 3.2. Tuatara Cloacal Swab Sample Collection

A total of 60 tuatara located on Takapourewa (Stephens Island), New Zealand (40.6702° S, 173.9969° E) were sampled during February and March 2021. Tuatara were caught by hand and held in dorsal recumbency for cloacal swab sampling. Snout-vent lengths and weight measurements were also recorded. Cloacal swabs were placed into RNA stabilisation solution (RNAProtect Tissue Reagent, Qiagen) and stored at 4°C for up to one month while on Takapourewa. On returning to the mainland, swabs were placed in a −80°C freezer until RNA was extracted.

### 3.3. Tuatara Cloacal Swab Total RNA Extraction

Frozen cloacal swabs were defrosted before being placed in ZR BashingBead Lysis Tubes (0.1 and 0.5mm) (Zymo Research) filled with 1mL of DNA/RNA shield (Zymo Research). Lysis tubes were placed into a mini-beadbeater 24 disruptor (Biospec Products Inc.) and were homogenised for five minutes. RNA was extracted using the ZymoBIOMICS MagBead RNA kit (Zymo Research) with a few additions to the manufacturers protocol. Briefly, three additional molecular grade pure ethanol washes were undertaken to remove any residual guanidine contamination. RNA was quantified using a nanodrop. RNA from 55 of the 60 cloacal swabs were at suitable concentrations. Equal concentrations of RNA from 6-8 individuals were pooled into eight groups based on the BCI of the individuals, which is a measure of overall health of the tuatara (BCI= log weight / log snout vent length). BCI ranged from an average of 1.025 (T1) to 1.197 (T8) (Table S1) (12).

### 3.4. Tuatara RNA Sequencing

Extracted RNA was subject to total RNA sequencing. Libraries were prepared using the Illumina Stranded Total RNA Prep with Ribo-Zero Plus (Illumina) and 16 cycles of PCR. Paired-end 150bp sequencing of the RNA libraries was performed on the Illumina NovaSeq 6000 platform.

### 3.5. Virome Assembly and Virus Identification

Paired reads were trimmed and assembled *de novo* using Trinity v2.11 with the “trimmomatic” flag option (13). Sequence similarity searches against the NCBI nucleotide (nt) database and the non-redundant (nr) protein database using BLASTn and Diamond (BLASTx), respectively, were used to annotate assembled contigs (14). A maximum expected value of 1×10^5^ was used as a cut-off to filter putative viral contigs. Contigs were categorised into higher kingdoms using the BLASTn “sskingdoms” flag option. Non-viral blast hits including host contigs with sequence similarity to viral transcripts (e.g. endogenous viral elements) were removed from further analysis. Viral contigs that have previously been identified as viral contaminants from laboratory components were also removed from further analysis (15). Based on the BLASTn and Diamond results, putative viral contigs were further analysed using Geneious Prime 2020.2.4 to find and translate open reading frames (ORFs).

### 3.6. Protein Structure Homology Searching for Viral Discovery

We used a protein structure homology search to identify highly divergent viral transcripts that did not share significant sequence similarity to other known transcripts. Such “orphan contigs” (16) were translated into ORFs using the EMBOSS getorf program (17), with the minimum nucleotide size of the ORF set to 1,000 nucleotides, the maximum nucleotide size of the ORF set to 50,000 and the “methionine” flag option set to only report ORFs with a methionine amino acid starting codon. Reported ORFs were submitted to Phyre2, which uses remote homology detection to build 3D models to predict and analyse protein structure and function (18). Predicted Protein Data Bank (PDB) protein structures were aligned with PDB templates and visualised using ChimeraX-1.3 (19,20), with default settings. Virus sequences with predicted polymerase structures with a confidence value of 90% were aligned with representative protein sequences from the same viral family or order obtained from NCBI RefSeq using MAFFT v7.450 (E-INS-I algorithm). Conserved domains were visually confirmed before phylogenetic trees were estimated using the same method outlined in section 3.8.

### 3.7. Viral Abundance Estimations

Viral abundance was estimated using Trinity’s “align and estimate” tool. RNA-Seq by Expectation-Maximization (RSEM) (21) was selected as the method of abundance estimation, Bowtie2 (22) as the alignment method and the “prep reference” flag enabled. To mitigate the impact of contamination due to index-hopping, viral transcripts with expected abundances of less than 0.1% of the highest expected abundance for that virus across other libraries were removed from further analysis. Total viral abundance estimates associated with eukaryotic hosts (excluding those from plant, fungi and protist hosts) and/or taxonomic orders across all libraries were compiled. Estimated abundances were standardised to the number of paired reads per library and normalised by dividing by the total standardised viral abundances within a virus family/order across all of the libraries.

### 3.8. Virus Phylogenetic Analysis

Translated viral protein polymerase sequences (i.e. RNA-dependent RNA polymerase, RdRp) were aligned with representative protein sequences from the same taxonomic viral family or order obtained from NCBI RefSeq using MAFFT v7.450 (L-INS-I algorithm or E-INS-I algorithm) (alignment lengths denoted in Table S2) (23). Poorly aligned regions were eliminated using trimAL v1.2 with the gap threshold flag set to 0.9 (24). IQ-TREE v1.6.12 was used to estimate maximum likelihood phylogenetic trees for each viral family/order (25). The LG amino acid substitution model was selected with 1000 ultra-fast bootstrapping replicates and the -alrt flag specifying 1000 bootstrap replicates for the SH-like approximate likelihood ratio test. Phylogenetic trees were annotated using the Interactive Tree of Life v6.5.2 (26).

### 3.10 Viral Nomenclature

In cases where viruses fell into known species, we have provided a tentative virus species name (prior to formal verification by the International Committee on Taxonomy of Viruses (ICTV)). A virus was arbitrarily considered a novel species if it shared <90% amino acid similarity within the most conserved region (i.e. RdRp/polymerase) (27). For novel virus sequences we have provided a proposed virus name.

## 4. Results

Cloacal swabs were sampled from tuatara located on Takapourewa, New Zealand. Total RNA derived from these were pooled based on BCI and were sequenced (Figure 1a). Metatranscriptomic sequencing of cloacal swab samples yielded sequencing libraries of between 44-58 million paired-end reads that were assembled into 362,973–756,621 contigs. A large proportion (71-84%) of the standardised abundance estimates across each of the libraries were classified as unassigned, while 0.02-0.03% were classified as viral (Figure 1b).

**Figure 1.**
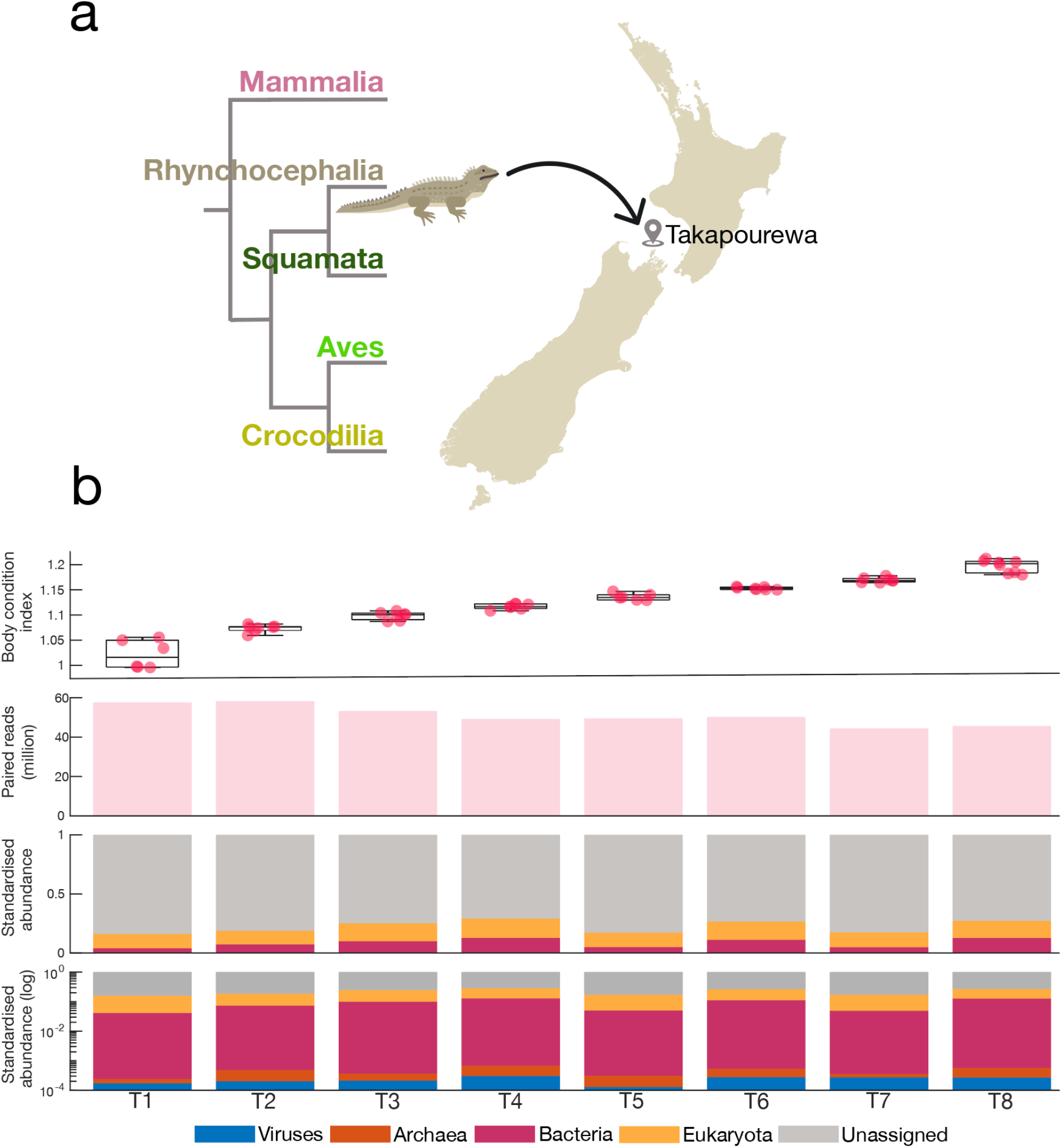
**(a)** Phylogenetic position of the order Rhynchocephalia and a map indicating the sampling location of tuatara from Takapourewa (Stephens Island), New Zealand. Tuatara illustration by Hamish Thompson. (**b**) (top panel) BCI measures of individual tuatara across each library, T1-T8. (second panel) Total paired-end sequencing reads from cloacal swab metatranscriptome libraries. (third panel) Standardised and normalised abundance estimates of viral, archaeal, bacterial, eukaryotic and unassigned transcripts identified in cloacal swab metatranscriptome libraries. (bottom panel) Logarithmic standardised and normalised abundance estimates of viral, archaeal, bacterial, eukaryotic and unassigned transcripts identified in cloacal swab metatranscriptome libraries.

### 4.1 Viral Abundance and Diversity

Analysis of the tuatara cloacal metatranscriptome libraries revealed that the viromes of tuatara are highly diverse (Figure 2). Exogenous viral transcripts from 42 viral families/orders that are genetically related to viruses that have previously been known to infect vertebrates, invertebrates, bacteria, plants, fungi and protists, were identified across the tuatara cloacal metatranscriptome.

**Figure 2.**
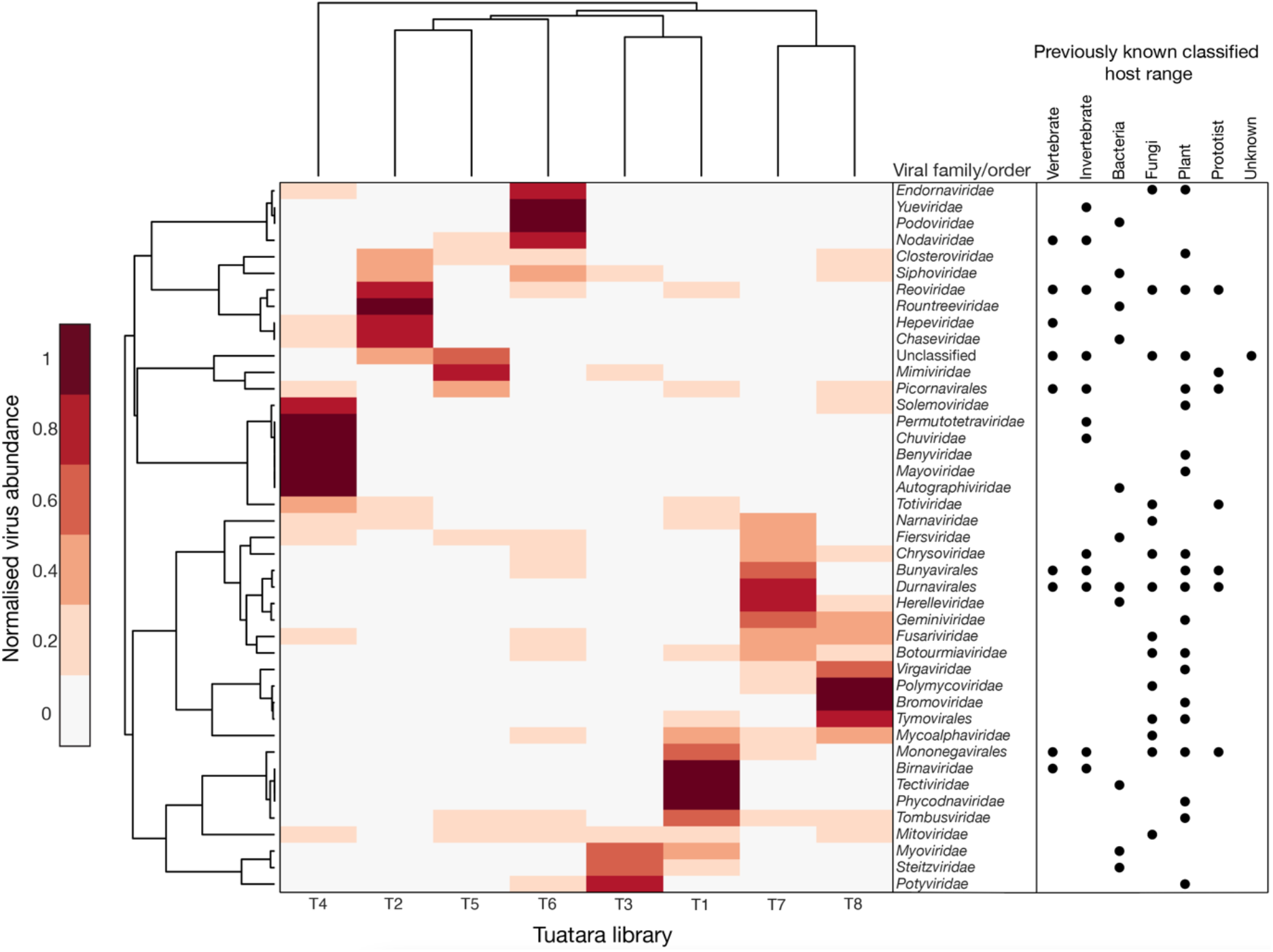
Clustergram of normalised viral transcript abundance grouped by viral family or order identified in the tuatara cloacal metatranscriptome libraries. Information regarding viral family and order level host ranges were obtained from the ICTV and through a literature search.

We focused our analysis on viral transcripts likely infecting eukaryotic hosts excluding those associated with plants, fungi and protist hosts. This resulted in the identification of viral transcripts from 19 different viral families or orders from the eight tuatara BCI sequencing libraries (Figure 3). Overall viral richness ranged from six to 13 virus families or orders across the eight libraries. We found that all sequencing libraries contained transcripts belonging to the *Virgaviridae, Picornavirales* and *Durnavirales* (Figure 3).

**Figure 3.**
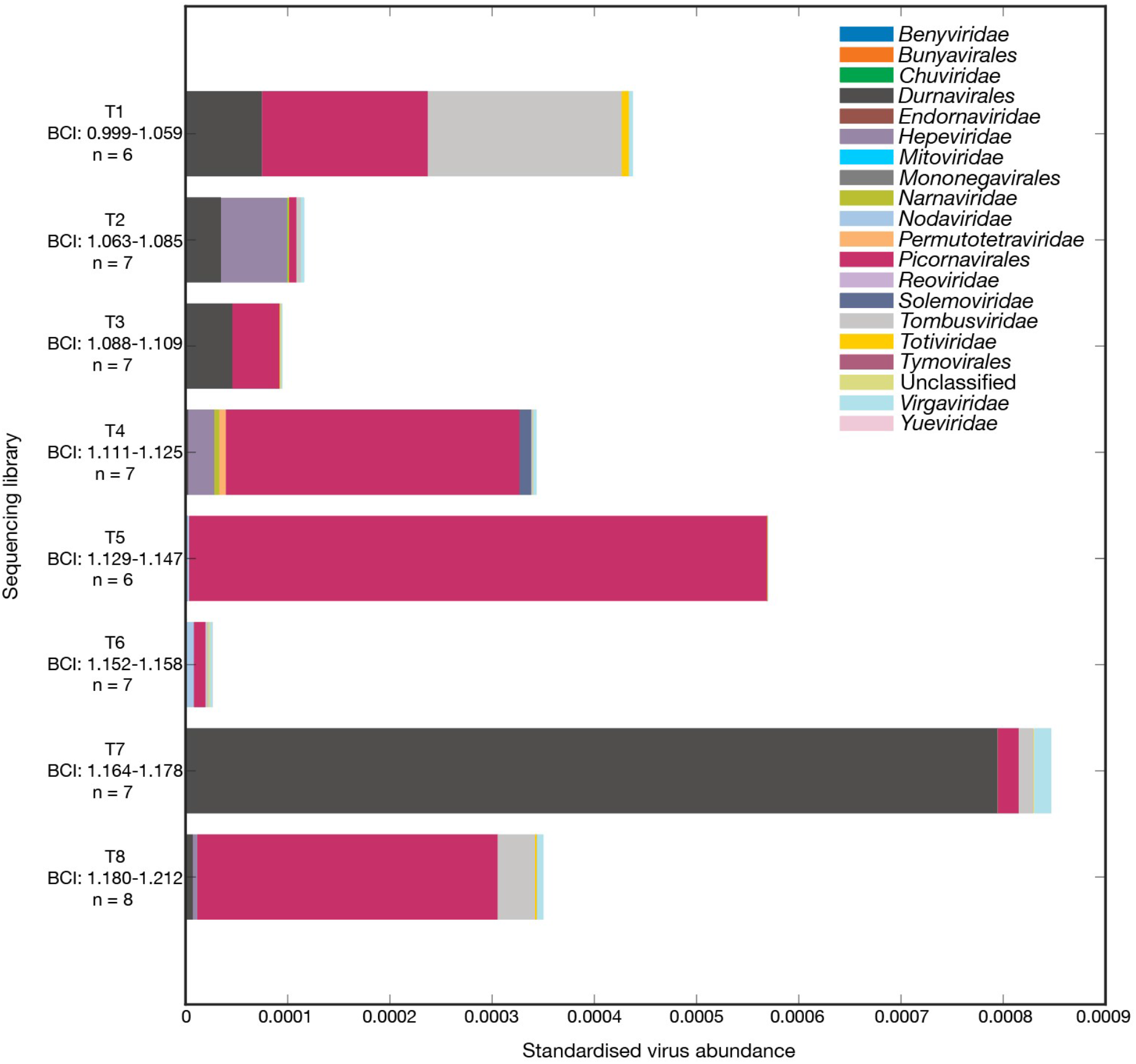
Bar graph of standardised abundance of eukaryotic (excluding plants, fungi and protist) host associated viral transcripts coloured by viral family or order. Host information was inferred from the closest genetic relative identified through a BLAST search. Ranges of individual BCI measurements per sequencing library and the number of individuals that were pooled per sequencing library is reported.

### 4.2 Novel Tuatara Cloaca-Associated Viruses

Of the viral transcripts identified in the tuatara cloacal metatranscriptomes a number contained partial or complete RdRp transcripts that allowed the identification of 56 exogenous viruses, 48 of which were potentially novel dietary related species, and one highly divergent novel *Picornaviridae* species that is likely associated with a vertebrate host (see Table S2 for contig lengths). We refer to these as ‘tuatara cloaca-associated’ viral transcripts since it was challenging to establish the true host of these viruses as the majority were likely either dietary related or exhibited a large degree of diversity. In addition, many of their closest genetic relatives found in public databases had limited or misassigned taxonomic information regarding host species (28). In addition, we identified two distinct novel endogenous tuatara adintoviruses. Due to the high proportion of seemingly dietary-related viruses determined by phylogenetic analysis, that would not actively replicate in the tuatara, it was not possible to investigate whether BCI significantly impacted beta diversity or viral composition as these viruses likely did not infect the tuatara. We now describe each viral taxa in turn.

#### 4.2.1 Mononegavirales

Three negative stranded RNA viruses from the order *Mononegavirales* were identified in the tuatara cloacal metatranscriptomes (Figure 4a). One of these, provisionally termed Tuatara cloaca-associated rabdovirus-1 (standardised abundance 9.01×10^-8^), was further classified as belonging to the family *Rhabdoviridae* as this virus fell alongside other rhabdoviruses sampled from both invertebrate and vertebrate hosts. The other two novel *Mononegavirales* were further classified as belonging to the *Mymonaviridae* and named Tuatara cloaca-associated mymonavirus-1 (standardised abundance 5.91×10^-7^) and Tuatara cloaca-associated mymonavirus-2 (standardised abundance 2.43×10^-7^). Both novel tuatara cloaca-associated mymonaviruses grouped with other mymonaviruses sampled from plant, fungi and arthropod hosts and were therefore likely dietary related viruses.

**Figure 4.**
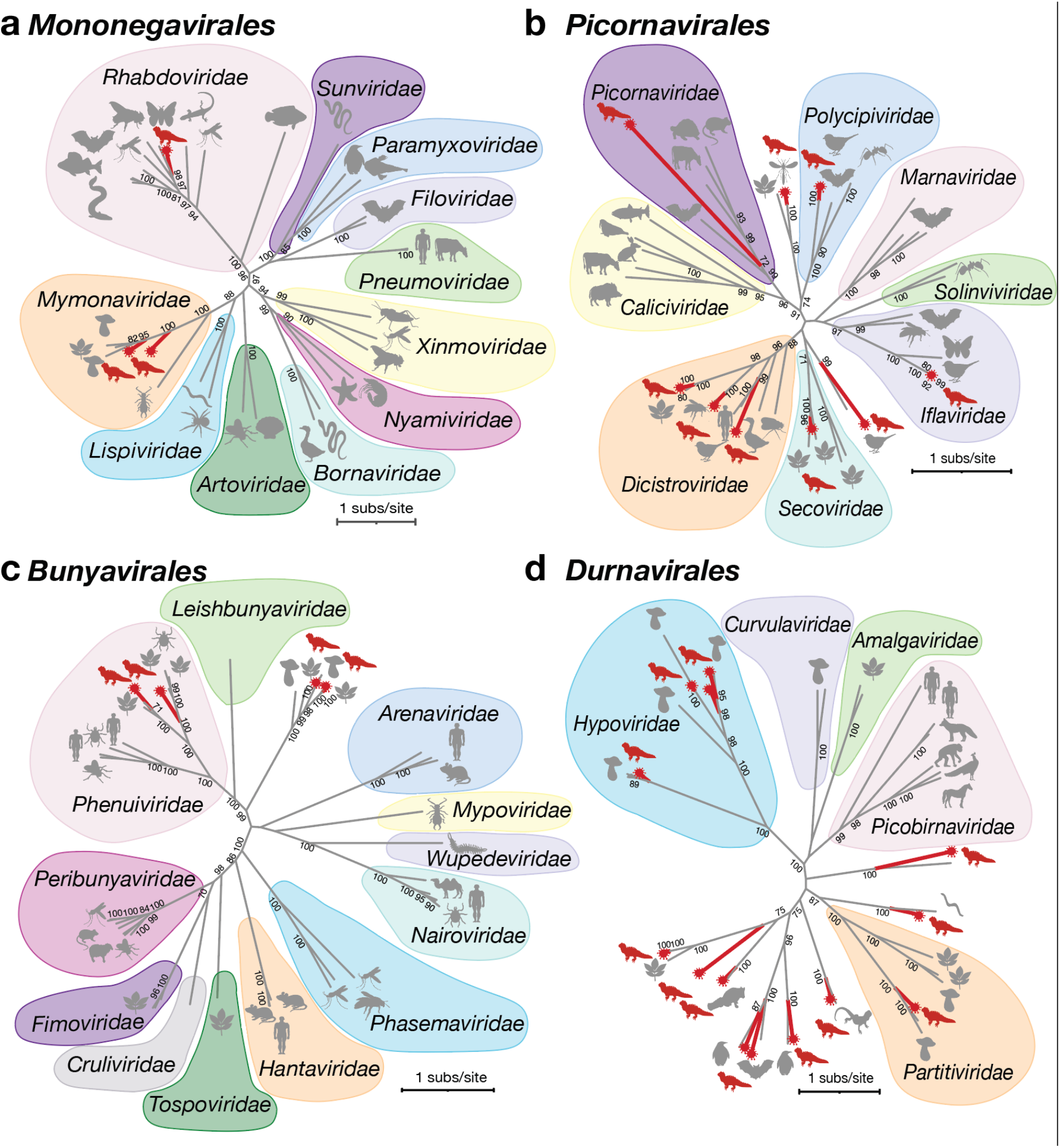
Maximum likelihood phylogenetic trees of representative viral transcripts containing RdRp from the orders (**a**) *Mononegavirales,* (**b**) *Picornavirales,* (**c**) *Bunyavirales* and (**d**) *Durnavirales.* Tuatara cloaca-associated viruses are highlighted in red. Viruses classified to the family level are highlighted. Branches are scaled to the number of amino acid substitutions per site. All phylogenetic trees are unrooted. Nodes with bootstrap values of >70% are noted. Amino acid length of tuatara cloaca-associated transcripts identified in this study used for alignment can be found in Table S2.

#### 4.2.2 Picornavirales

We identified nine single-stranded positive-sense RNA viruses within the *Picornavirales,*eight of which were potentially novel, with one other virus – *Hubei picorna-like virus 51* - previously identified as infecting flying insects (order Odonata) (Figure 4b, Table S2). Of the eight potentially novel viruses, three were further classified as belonging to the *Dicistroviridae,* while others fell within the *Iflaviridae, Picornaviridae, Polycipiviridae, Secoviridae.* One virus could not be further classified to family-level and was named Tuatara cloaca-associated picorna-like virus-1. This virus was the most abundant virus identified (standardised abundance 1.46–10^-4^) and was closely related to a *Picornavirales* species that was identified in the metagenome of a bird and is therefore likely dietary related (Table S2). Notably, using a protein structural homology search we also identified a highly divergent member of the *Picornaviridae* provisionally termed Tuatara cloaca-associated picornavirus-1. This virus is described in more detail in section 4.3. The other tuatara viruses shared significant amino acid sequence similarity with viruses sampled from avian hosts, albeit via metagenome studies, as well as invertebrate and plant hosts, and are therefore likely dietary related (Table S2).

#### 4.2.3 Bunyavirales

Two potentially novel RNA viruses from the *Phenuiviridae* were identified, while a further two novel bunya-like viruses were also identified but could not be further classified to the family level (Figure 4c). All four potentially novel viral transcripts were most closely related to viruses sampled from plant or fungi hosts and were therefore unlikely to infect the tuatara (Table S2).

#### 4.2.4 Durnavirales

We identified 14 viruses within the *Durnavirales,* of which four belonged to the *Hypoviridae,* one to the *Partitiviridae,* and nine viral transcripts could not be categorised beyond order level (Figure 4d). All four tuatara cloaca-associated hypoviruses were closely related to fungi infecting hypoviruses, indicating that their true hosts are likely to lie within the fungi kingdom. Similarly, the virus that shared the closest amino acid sequence similarity to Tuatara cloaca-associated patitivirus-1(standardised abundance 1.91×10^-5^) was also a fungi associated virus (Table S2). The remaining nine viruses were closely related to vertebrate metagenome associated viruses as well as to environmental, plant and invertebrate associated viruses so that their true hosts are difficult to identify, but again are likely to be dietary related.

#### 4.2.5 Double-Stranded RNA Viral Transcripts

We identified two potentially novel viral species falling within the *Reoviridae* termed Tuatara cloaca-associated reovirus-1 and Tuatara cloaca-associated reovirus-2 with similar standardised abundances – 6.95×10^-8^ and 3.98×10^-8^, respectively (Figure 5a). The top BLAST result to both viral transcripts was *Bercke-Baary Melophagus reo-like virus* (UJG27935.1) associated with invertebrates, with 83.33% and 84.21% amino acid sequence identity, respectively (Table S2). Consequently, it is likely that the novel tuatara cloaca-associated reoviruses may be dietary related. The genera of the two novel tuatara viruses within the *Reoviridae* is uncertain as they fell among unclassified reoviruses that formed a sister clade to the *Fijivirus* genus that infect plants.

**Figure 5.**
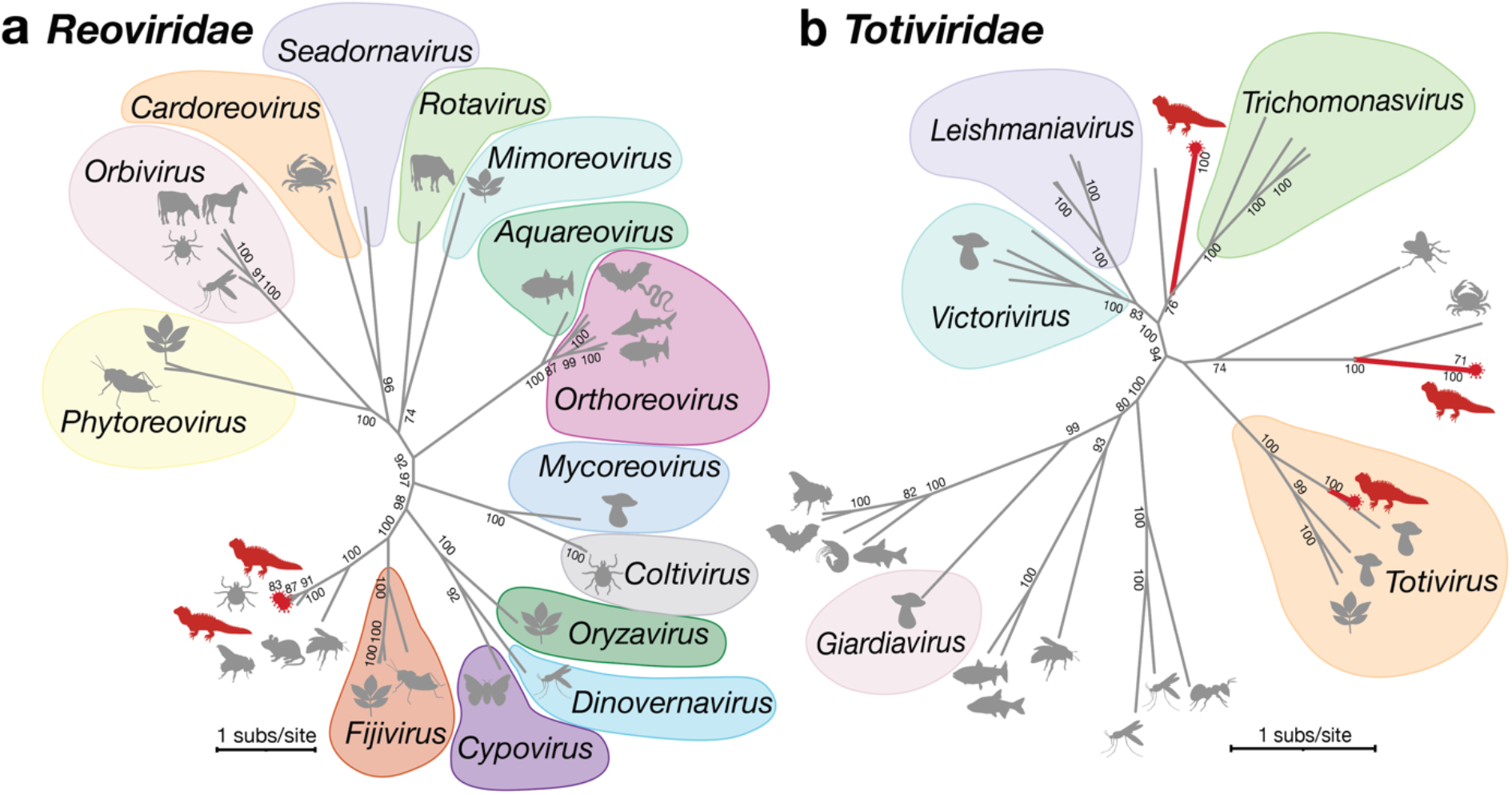
Maximum likelihood phylogenetic trees of representative double-stranded (ds) RNA viral transcripts containing RdRp from the (**a**) *Reoviridae* and (**b**) *Totiviridae.* Tuatara cloaca-associated viruses are highlighted in red. Viruses classified at the genus level are highlighted. Branches are scaled to the number of amino acid substitutions per site. All phylogenetic trees are unrooted. Nodes with bootstrap values of >70% are noted. Amino acid length of tuatara cloaca-associated transcripts identified in this study used for alignment can be found in Table S2.

Three novel tuatara cloaca-associated double-stranded RNA viruses within the *Totiviridae* were identified in the tuatara cloacal metatranscriptomes and termed Tuatara cloaca-associated totivirus-1, 2 and 3 (Figure 5b). Phylogenetic analysis of the Tuatara cloaca-associated totivirus-3 (standardised abundance 5.41×10^-7^) revealed that this novel virus fell within the genus *Totivirus* alongside the closest genetic relative, *Erysiphe necator associated totivirus 7,* which shared 61.9% amino acid identity (Table S2). Both Tuatara cloaca-associated totivirus-1 and 2 could not be classified beyond the family level. All three novel tuatara totiviruses are likely dietary related viruses as they were closely related to environmental, invertebrate and fungi host associated viruses.

#### 4.2.6 Single-Stranded RNA Viral Transcripts

We identified a single-stranded RNA virus in the *Chuviridae,* tentatively called Tuatara cloaca-associated chuvirus (standardised abundance 1.63×10^-7^) (Figure 6a). Phylogenetic analysis could not classify this transcript beyond family level although it formed an outgroup to *Hubei odonate virus 11* from the Odonata (flying insects) and is therefore likely dietary related.

**Figure 6.**
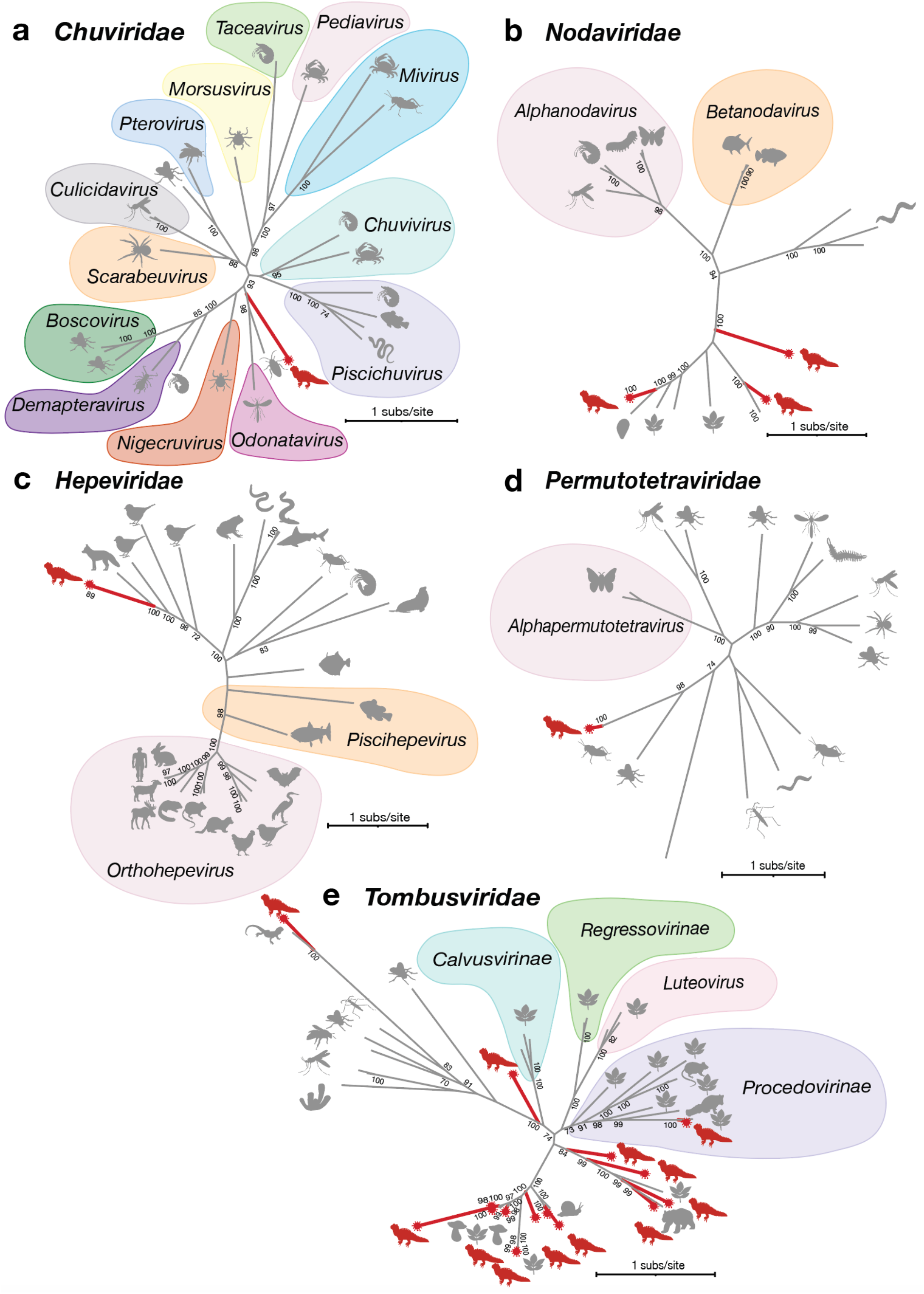
Maximum likelihood phylogenetic trees of representative single-stranded (ss) RNA viral transcripts containing RdRp from the (**a**) *Chuviridae,* (**b**) *Nodaviridae,* (**c**) *Hepeviridae,* (**d**) *Permutotetraviridae*, (**e**) *Tombusviridae*. Tuatara cloaca-associated viruses are highlighted in red. Viruses classified at the genus or subfamily level are highlighted. Branches are scaled to the number of amino acid substitutions per site. All phylogenetic trees are unrooted. Nodes with bootstrap values of >70% are noted. Amino acid length of tuatara cloaca-associated transcripts identified in this study used for alignment can be found in Table S2.

Three viruses were identified within the *Nodaviridae*, none of which could be classified beyond the family level as all three viruses fell close to viruses that have yet to be classified and likely infect invertebrate, plant and environmental hosts and are therefore, unlikely to infect the tuatara (Figure 6b).

A novel single-stranded positive-sense virus within the *Hepeviridae,* termed Tuatara cloaca-associated hepevirus-1 (standardised abundance 2.52×10^-5^ – 4.82×10^-7^), was uncovered in three of the eight tuatara cloacal metatranscriptome sequencing libraries, all sharing greater than 98.4% amino acid sequence identity (Figure 6c). This Tuatara cloaca-associated hepevirus was most closely related to *Swiper virus,* identified in the faecal metagenome of a red fox (*Vulpes vulpes*) and multiple avian viruses identified from cloacal metagenomes and are likely dietary related viruses (Table S2).

We identified one single-stranded positive-sense virus within the *Permutotetraviridae* in the tuatara cloacal metatranscriptome (Figure 6d). This virus shared >95% amino acid similarity with the *Hubei permutotetra-like virus 10* (e-value 2×10^-50^) (Table S2). *Hubei permutotetra-like virus* was first identified in the Orthoptera order of insects (29) again suggesting that it is dietary associated.

Finally, we identified 15 single-stranded positive-sense viruses within the *Tombusviridae* (Figure 6e). Of these, ten were classified as potentially novel tuatara cloaca-associated tombusviruses, while five had previously been discovered and classified (Table S2). One of the ten novel viruses Tuatara cloaca-associated tombusvirus-8 grouped with plant viruses and other viruses identified in metagenomic studies of vertebrates in the subfamily *Procedovirinae* and was therefore likely a dietary related virus (Table S2). While the remaining nine novel tuatara cloaca-associated tombusviruses did not fall within currently classified subfamilies of the *Tombusviridae*, they were also likely dietary related as they were phylogenetically similar to viruses identified in vertebrate metagenomic data as well as to viruses infecting invertebrates, fungi and plants (Table S2).

#### 4.2.7 Adintoviridae

We identified five double-stranded DNA adintoviral transcripts from two distinct novel adintoviruses in the tuatara cloacal metatranscriptomes. The standardised abundance of the Tuatara adintovirus-1 and Tuatara adintovirus-2 transcripts was 1.47×10^-6^-5.88×10^-7^ and 4.86×10^-6^, respectively. Due to their high degree of fragmentation these were likely endogenous viruses (Figure 7). Both novel tuatara adintoviruses fell within the *betadintovirus* genus and grouped with several reptilian adintoviruses including an endogenised *Crocodylus saltwater crocodile adintovirus* and an exogenous *Terrapene box turtle adintovirus,* as well as exogenous and endogenous fish adintoviruses. Interestingly, the phylogeny of the betadintoviurses mirrors that of the host phylogeny indicating that the adintoviruses have likely codiverged with the tuatara over time.

**Figure 7.**
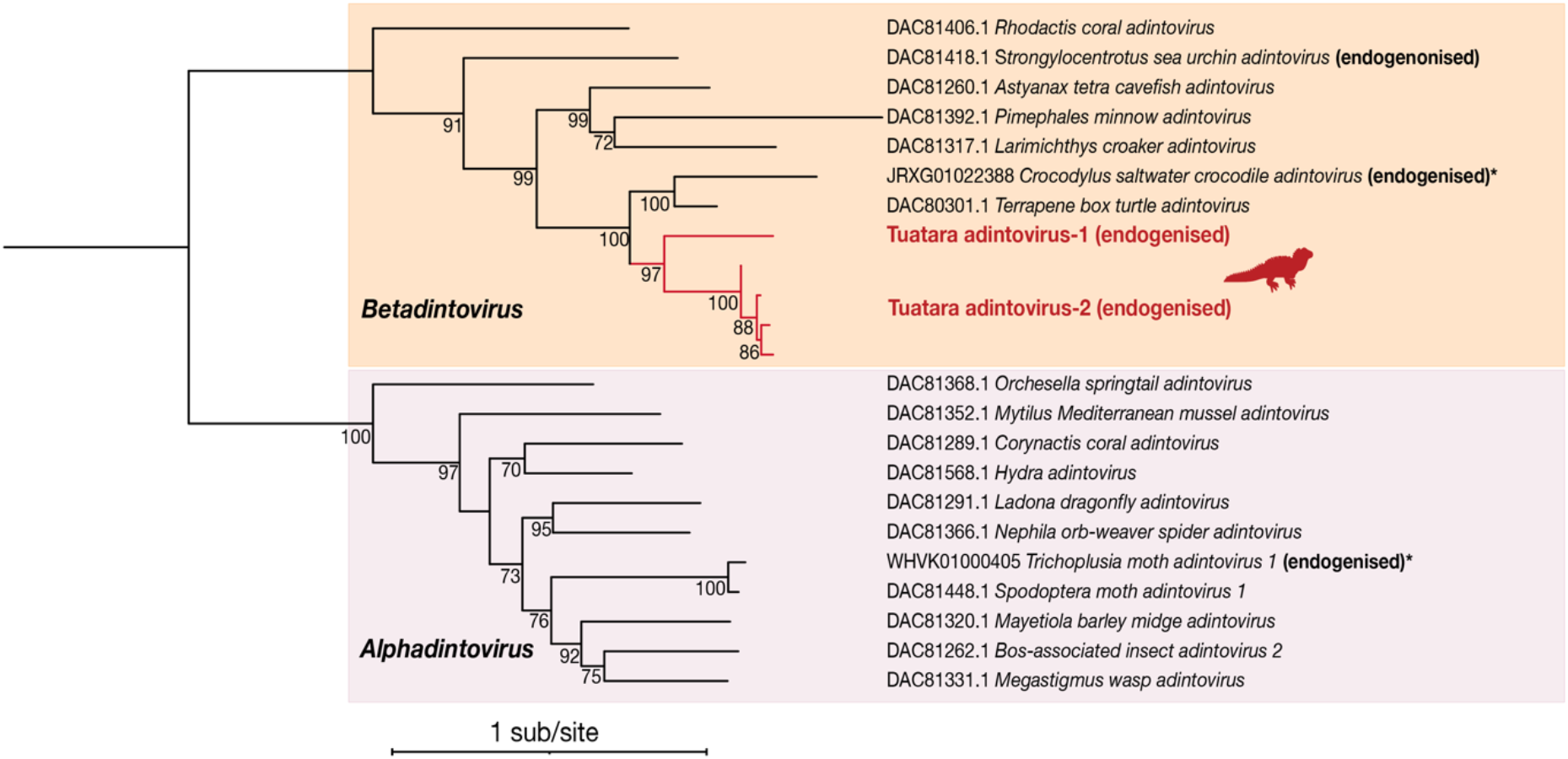
Maximum likelihood phylogenetic tree of representative double-stranded (ds) DNA viral transcripts within the *Adintoviridae* containing PolB. Endogenised tuatara adintoviruses are highlighted in red. Branches are scaled to the number of amino acid substitutions per site. Phylogenetic tree is rooted at the midpoint. Nodes with bootstrap values of >70% are noted. Amino acid length of tuatara cloaca-associated transcripts identified in this study used for alignment can be found in Table S2. Sequences denoted with an asterisk cannot be found on NCBI and were taken from Starrett *et al.,* 2021 (30).

### 4.3 Identifying Viruses Through Protein Structural Homology

We used a protein structure homology based search to identify highly divergent novel viruses from the contigs that did not share significant sequence similarity to other known sequences. We identified a total of 2,900,558 such orphan contigs within the eight tuatara cloacal metatranscriptomes, of which 1,430 contained an ORF of >1000 nucleotides in length. These were submitted to Phyre2 for structural similarity analysis which detected five putative viral structures with a confidence of >90% (18) (Figure 8). The top structural hit, from a 485aa orphan contig identified in the T7 library, was to a RdRp structure from *Rabbit hemorrhagic disease virus* (RHDV) (Protein Data Bank identifier 1KHV), a member of the *Caliciviridae* (order *Picornavirales),* with a confidence of 91.5% and 27% percentage identity (Figure 9a).

**Figure 8.**
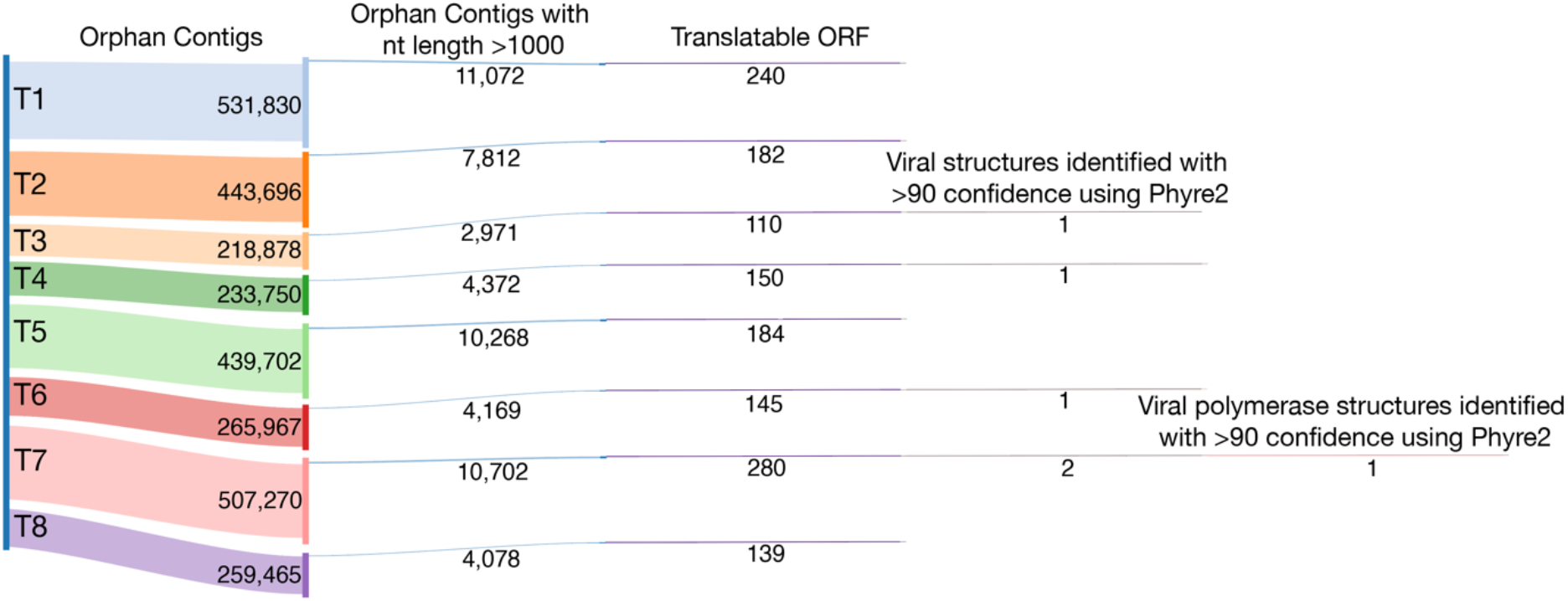
Sankey diagram depicting the stages of virus discovery using a protein structural homology search based approach. The sankey diagram was made using SankeyMATIC.

**Figure 9.**
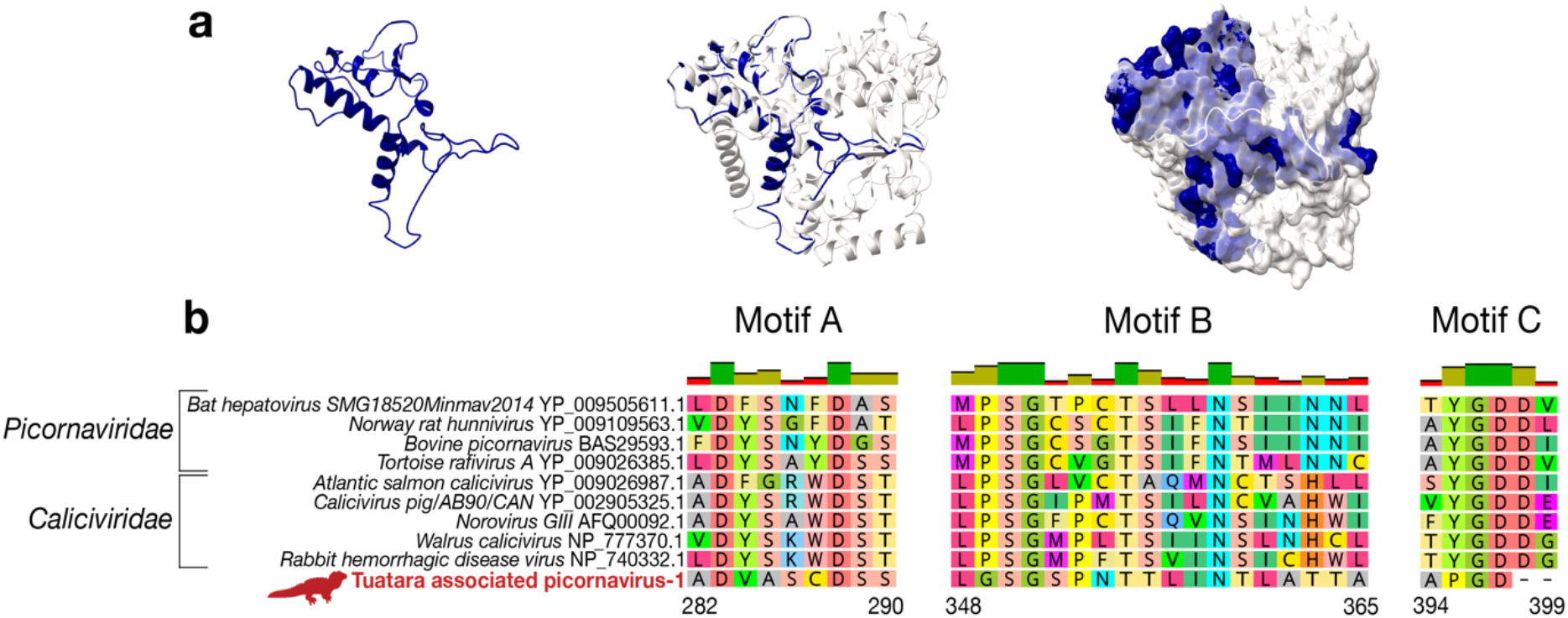
Protein structural prediction and conserved amino acid motif analysis of the highly divergent tuatara cloaca-associated picornavirus-1. (**a**) (left) A 3D model prediction of the tuatara cloaca-associated picornavirus-1 protein sequence coloured blue. (middle) Ribbon structural alignment of the 3D predicted model of the tuatara cloaca-associated picornavirus-1 protein sequence coloured blue and the RdRp structure of RHDV coloured white. (right) Molecular surface structural alignment of the 3D predicted model of the tuatara cloaca-associated picornavirus-1 protein sequence coloured blue and the RdRp structure of RHDV coloured white. (**b**) Alignment of multiple viral amino acid sequences from the RdRp showing three domains (Domain IV-VII) conserved across members of the *Picornaviridae* and *Caliciviridae*, including the novel and highly divergent Tuatara cloaca-associated picornavirus-1 (highlighted in red).

Phylogenetic analysis of this novel tuatara viral sequence and representative RdRp sequences from across the *Picornavirales* confirmed that the contig contained the core A, B and C RdRp motifs of the *Picornavirales* (31,32) (Figure 9b). Phylogenetic analysis revealed that the novel tuatara viral sequence fell between the genera *Bopivirus* and *Hepatovirus* within the *Picornaviridae* (Figure 4b). Notably, because the novel tuatara viral sequence clustered with viruses sampled from other vertebrate hosts including bats, cattle, tortoises and rats, albeit highly divergent, this virus transcript may indeed infect the tuatara. We have tentatively named this highly divergent tuatara virus, Tuatara cloaca-associated picornavirus-1 (standardised abundance 3.06×10^-6^) (Table S2).

## 5. Discussion

We analysed the viromes of tuatara located on Takapourewa, a remote off-shore island in New Zealand that is home to the largest naturally occurring population of tuatara (>30,000 individuals) (1). Exogenous viral transcripts spanning 42 viral families/orders were identified. These included 49 novel exogenous RNA tuatara cloaca-associated viruses, along with a further seven already classified virus species. Of the 49 novel viruses discovered here, 48 were likely dietary-related while one novel highly divergent virus within the *Picornaviridae,*Tuatara cloaca-associated picornavirus-1, may be a virus specific to tuatara. Similar findings were obtained in viral transcripts recently identified in the faecal virome of *Phrynocephalus* lizards (33) with considerable overlap with the tuatara virome described here, including viruses across 19 viral families shared among both viromes.

A further two endogenous adintoviruses were identified that phylogenetically grouped with other previously identified reptilian adintoviruses suggesting that, prior to endogenisation, adintoviruses may have once infected tuatara. The *Adintoviridae* is the only family within the *Orthopolintovirales* order, which was only formally classified in 2021 (34). To date, adintoviruses have either been discovered as whole or fragmented sequences endogenised into host genomes, or as exogenous DNA viruses (30). While two distinct endogenous tuatara adintoviruses were identified in the cloacal metatranscriptome it is likely that they originated from a single exogenous adintovirus that became endogenised into the tuatara genome and subsequently acquired a number of mutations (35). The discovery of these endogenous viral sequences, in addition to the >450 endogenous retroviruses that have previously been described (3–6), adds to our understanding of the evolutionary history of viral-tuatara codivergence.

While viruses do not contain a universally conserved gene like fungi and bacteria, the RdRp is characteristic of RNA viruses (36,37), playing a critical role in replicating the viral genome and carrying out transcription (37). Importantly, the core structural features of the RdRp are conserved despite considerable differences at the amino acid level (37) and, therefore, protein structure homology searches can be used to successfully identify highly divergent novel viruses (16). Using this approach, we identified a highly divergent novel tuatara cloaca-associated virus within the *Picornaviridae* termed Tuatara cloaca-associated picornavirus-1. Despite being highly divergent, the Tuatara cloaca-associated picornavirus-1 maintained the conserved RdRp motifs A-C (31,32). This novel virus likely infects vertebrates as it clustered with other *Picornaviridae* vertebrate infecting viruses sampled from bats, cattle, tortoises and rats. Picornaviruses are a family of highly diverse viruses that have been found to infect all classes of vertebrates (38). While the vast majority of picornavirus infections are asymptomatic they can cause disease in humans and livestock (38), and a picornavirus was recently found to cause kidney disease and shell weakness syndrome in European tortoise species (39). The discovery of the novel highly divergent Tuatara cloaca-associated picornavirus-1 expands our understanding of the highly diverse *Picornaviridae* family and highlights the utility of using a structural based approach to identify highly divergent novel viruses.

A major limitation of metatranscriptomics is that the true host range of many viruses often cannot be determined. Cloacal swabs and faecal samples are particularly challenging to identify viral-host associations as a high proportion of the total viral sequences identified are often dietary-related (33,40–42). Indeed, 48 of the 49 novel viruses identified in this study were likely dietary-related and, consequently, the hosts of these viral sequences remain unclear. Takapourewa tuatara primarily consume an insectivorous diet but are also known to eat fairy prion (*Pachyptila turtur*), fluttering shearwater (*Puffinus gavia*) and sparrow (*Passer domesticus*) chicks and eggs and kawakawa fruit (*Piper excelsum*) (1). Plant material and soil have also previously been identified in tuatara scats found on Takapourewa (1). Additionally, typical misassignment or limited taxonomic information regarding hosts of viral sequences found in public databases also makes it increasingly difficult to reveal viral-host relationships (28). While the novel Tuatara cloaca-associated picornavius-1 identified in this study likely infects vertebrates and may infect the tuatara, the true host remains unclear since tuatara consume vertebrates (1).

The tuatara population on Takapourewa is a natural, relict population that has been isolated for thousands of years (1). Unlike previous studies (33,40–42), an amino acid similarity approach was unable to identify tuatara nor vertebrate-specific viruses within cloacal samples. Strikingly, over 2.9 million orphan contigs were identified contributing to 65% of the total contigs assembled in the metatranscriptome data. As the Takapourewa tuatara population has been isolated for thousands of years it is likely that a high proportion of the orphan contigs are indeed viral and highly divergent but due to current computational limitations that rely heavily on sequence or structure homology, they are unable to be identified.

The discovery of 49 potentially novel viruses emphasises that we are only just beginning to explore the virosphere, with our reliance on identifying viruses that share significant sequence or structural similarity to known viruses found in publicly available databases a fundamental challenge of discovering novel viruses. Indeed, there are an estimated 1×10^31^ viruses present on Earth (43), yet, fewer than 10,500 of these viruses have been formally classified (34), in part due to a historic sampling bias that has favoured the sampling and sequencing of economically and medically relevant viral species (28,44,45). Even more so, there is a very limited representation of viral structures within databases. For example, the majority (66%) of the viral structures in the PDB are structures resolved solely from human viruses (28). This study emphasises the importance of sequencing the viromes of neglected and under-sampled species such as the tuatara to further expand our knowledge of known viruses which will consequently enable us to identify a broader range of novel viral sequences.

## 6. Acknowledgements

We would like to acknowledge Professor Nicola J. Nelson, Victoria University of Wellington, and Noela McGregor, Ngāti Kōata Trust, for supporting our research on discovering viruses in tuatara found on Takapourewa, and Laura Rutten, Susan Keall, Diane Ormsby, and Joseph Altobelli for helping assist with sampling and field work.

S.J.W. is supported by a University of Otago Doctoral Scholarship, J.L.G. is funded by a New Zealand Royal Society Rutherford Discovery Fellowship (RDF-20-UOO-007) and E.C.H. is funded by an ARC Australian Laureate Fellowship (FL170100022). This project was also supported by the University of Otago School of Biomedical Sciences Dean’s Bequest Fund Grant.

## 7. Supplementary Material

**Table S1.** BCI calculations used to pool individual tuatara cloacal swab RNA into eight libraries

**Table S2.** Tuatara cloaca-associated viruses identified in the cloacal metatranscriptome libraries and the top viral polymerase containing amino acid sequence hits from the NCBI nr database.

## 8. Data Availability

Raw reads are available via [pending] while virus sequences are available on GenBank under accession numbers [pending].

